# Promotion of new expression of connexin gene Cx46 (*GJA3*) in the cochlea after Cx26 (*GJB2*) deficiency

**DOI:** 10.1101/2025.03.30.646179

**Authors:** Tian-Ying Zhai, Jin Chen, Yong Kong, Chun Liang, Hong-Bo Zhao

## Abstract

Connexin 26 (Cx26, *GJB2*) mutations induce a high incidence of hearing loss, responsible for 70-80% of nonsyndromic hearing loss. The pathological changes mainly locate in the cochlea. However, the genetic changes in the cochlea after deficiency of Cx26 remail unclear, which hampers to fully understand the underlying deafness mechanisms to develop therapeutic interventions. In this study, we employed bulk Poly(A) RNA-Seq technique and found that Cx26 deficiency could cause many genes up-regulating and down-regulating in the cochlea. A significant change was that Cx46 (*GJA3*), which is like Cx26 but expresses in the eye rather than the ear normally, had a remarkable upregulation and occurred in the cochlea after Cx26 deficiency. Immunofluorescent staining confirmed that Cx46 had expression in the cochlea and integrated into the gap junction networks among the cochlear supporting cells and in the cochlear lateral wall at the same location as Cx26 expression. Moreover, newly expressed Cx46 could be found in the same gap junctional plaques with Cx26. In addition, this promotion of new Cx46 expression is Cx26-specific; there was no promotion of Cx46 expression in the cochlea after deletion of Cx30 (*GJB6*), which also predominantly co-expresses with Cx26 in the cochlea. These data demonstrated that Cx26 deficiency could promote Cx26-like Cx46 expression in the cochlea for compensation. This finding also provides a new cue for developing a genetic approach to treat this common hereditary deafness induced by *GJB2* mutations.

**Highlight:**

- New Cx46 compensatively expresses in the cochlea after Cx26 deficiency
- Cx46 expression occurs in the same location as Cx26 in the cochlea
- Cx46 promotion is Cx26-specific, no expression in the Cx30 KO cochlea

## Introduction

Gap junction (GJ) gene *GJB2* (Cx26) mutations cause >50% of nonsyndromic hearing loss in the clinic (1-3). The pathological changes of Cx26 deafness mutations and deficiency have been extensively investigated (4). The major pathological changes locate in the cochlea, including disorders of the cochlear development (5-7), degeneration of hair cells and other cochlear cells (6, 7), reduction of endocochlear potential (EP) generation (5, 8), and impairment of active cochlear amplification (9, 10). However, little is known about the genetic changes in the cochlea after Cx26 deficiency, which hampers to fully understand the underlying deafness mechanisms to develop genetic therapeutic interventions, which currently is not available.

GJ is an intercellular channel, docked by two hemichannels from two neighbor cells, and extensively exists in almost all organs and tissues (11). GJ proteins in vertebrates are mainly encoded by a connexin gene family, which has about 20 connexin subtypes. Different connexins have different cell- or organ-specific distributions. In the cochlea, Cx26 and Cx30 (*GJB6*) are predominant connexin isoforms and co-express in the cochlear supporting cells and the cochlear lateral wall but not in the hair cells (12-14). Cx46 (*GJA3*) usually expresses in the eye but not in the cochlea (15, 16). In this study, we reported an unexpected finding that Cx46 expression is promoted in the cochlea after Cx26 deficiency. This may provide a cue for developing a new strategy in gene therapy for this common *GJB2* mutation-induced deafness.

## Materials and methods

### Cx26 conditional deletion mice and Cx30 knockout mice

Cx26 conditional deletion was generated by a Cre-FloxP technique. Pax2-Cre male mice (the Mutation Mouse Regional Center, Chapel Hill, NC) were crossed with Cx26^*loxP/loxP*^ mice (EM00245, European Mouse Mutant Archive) to create Pax2-Cx26 conditional knockout (cKO, Pax2-Cre^+^/Cx26^loxP(+/+)^) mice (6). Cx30 KO mice were purchased from European Mouse Mutant Archive (EM00323) (5). All experimental procedures were conducted in accordance with the policies of the Yale University Animal Care & Use Committee.

### RNA-Seq and quantitative RT-PCR

The cochlear was removed from the temporal bone and placed in 1.5ml tubes and quick-frozen in liquid nitrogen. RNAs were extracted by use of absolute RNA Nanoprep Kit (#400753, Agilent, Santa Clara, CA, United States) following manufactural instruction. The extracted RNA was tested for quality and concentration using NanoDrop™ One/OneC Microvolume UV-Vis Spectrophotometer (no. ND-ONE-W, Thermo Fisher Scientific Inc.), and sent to Yale Center for Genome Analysis (YCGA) for RNA sequencing.

For quantitative RT-PCR (qPCR), the extracted RNAs (500 ng) were reversely transcribed into DNA by using iScript cDNA Synthesis Kit (Cat. #1708891, Bio-Rad Laboratories). Real-time PCR was performed by QuantStudio 6 Pro (Thermo Fisher Scientific) with PowerUp SYBR Green Master Mix (A25742, Thermo Fisher Scientific) and following primers: Cx46-F: 5’- GGC AAC CAG GAG GAC CTG GG-3’; Cx46 -R: 5’- GGA AAG GCC ACA GGG TTT CCT GG-3’; The conditions for amplification were as follow: initial denaturation with two steps: 50 °C for 2min, then increase to 95 °C for 10 min; the amplification cycle: 95°C for 30s, 60°C for 30s, 72°C for 45s, totally 60 cycles. The melting curve was also measured by 95°C for 15s, 60°C for 1min, and 95°C for 1s. Each sample was repeated in triplicate and averaged. Universal 18S (AM1718, Thermo Fisher Scientific) was used as an internal control. Data was analyzed by Design &Analysis Software 2.8.0 (Thermo Fisher Scientific). The gene expression changes were calculated by standard curves and normalized to wild-type (WT) mice (17).

### Immunofluorescent staining

As we previously described (13, 14), the freshly isolated cochlea was fixed with 4% paraformaldehyde in 0.1 M PBS (pH 7.4) for 30 min. After decalcification, the cochlear sensory epithelium was micro-dissected. After washed with 0.1 M PBS for 3 times, the tissue was incubated in a blocking solution (10% goat serum and 1% BSA in the PBS) with 0.1% Triton X-100 for 30 min at room temperature. The tissue then was incubated with primary antibody monoclonal mouse anti-Cx26 (1:400, Cat# 33-5800) and polyclonal rabbit anti-Cx46 (1:200, Cat# RD236806, Thermo Fisher Scientific) in the blocking solution at 4°C overnight. After washing out the primary antibodies with PBS, the reaction to a 1:600 dilution of secondary Alexa Fluor® 488 or 568 conjugated antibodies (Thermo Fisher Scientific) in the blocking solution followed at room temperature for 1 hr. After completely washing out, the section was mounted with a fluorescence mounting medium (H-1000, Vector Lab, CA) and observed under a Nikon AX confocal microscope system (18).

### Data processing and statistical analysis

The numbers of recording mice in each experiment were indicated in the related figure. Data were plotted by SigmaPlot. Data were expressed as mean ± SEM. Statistical analyses were performed by SPSS v18.0 (SPSS Inc. Chicago, IL). One-way ANOVA with Bonferroni *post hoc* test was used. The threshold for significance was P= 0.05.

## Results

### Cx46 upregulation in the cochlea in Cx26 deficient mice

Fig. 1 shows upregulation of Cx46 expression in the cochlea after Cx26 deficiency in RNA-Seq examination. Volcano plot demonstrated that Cx46 expression was significantly upregulated in Cx26^+/-^ hetero-deletion mice (Fig.1A).

**Fig. 1.**
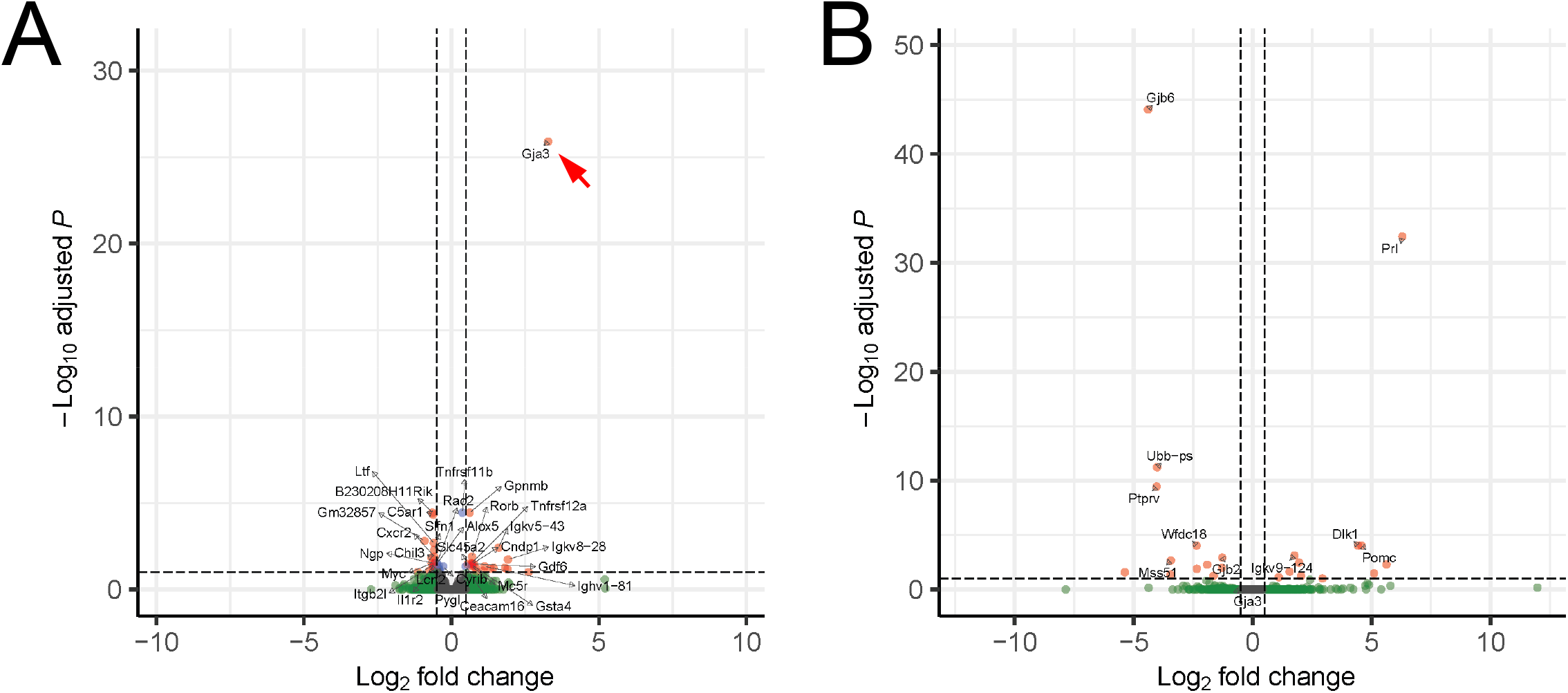
Cx46 (*GJA3*) upregulation in the cochlea after Cx26 (*GJB2*) deficiency but not Cx30 (*GJB6*) deficiency in RNA-Seq examination. Cx46 (*GJA3*) upregulation in the cochlea after Cx26 (*GJB2*) deficiency but not Cx30 (*GJB6*) deficiency in RNA-Seq examination. **A**: Volcano plot of gene upregulation and downregulation in the cochlea in Cx26 heterozygous knockout (KO) mice. A red arrow indicates significant upregulation of Cx46. **B:** No upregulation of Cx46 in the cochlea after Cx30 KO.

The promotion of Cx46 expression in the cochlea after Cx26 deficiency was further verified by real-time quantitative RT-PCR. Fig. 2 shows that Cx46 expression in the cochlea in Cx26 heterozygous and homozygous deletion (Cx26 Het and KO, respectively) mice were significantly increased. In comparison with WT mice, the levels of Cx46 expression in the cochlea in Cx26^+/-^ Het and Cx26^-/-^ KO mice were increased by 6.32±1.25 (n=5, P<0.05) and 6.6±1.01 (n=7, P<0.01) folds, respectively.

**Fig. 2.**
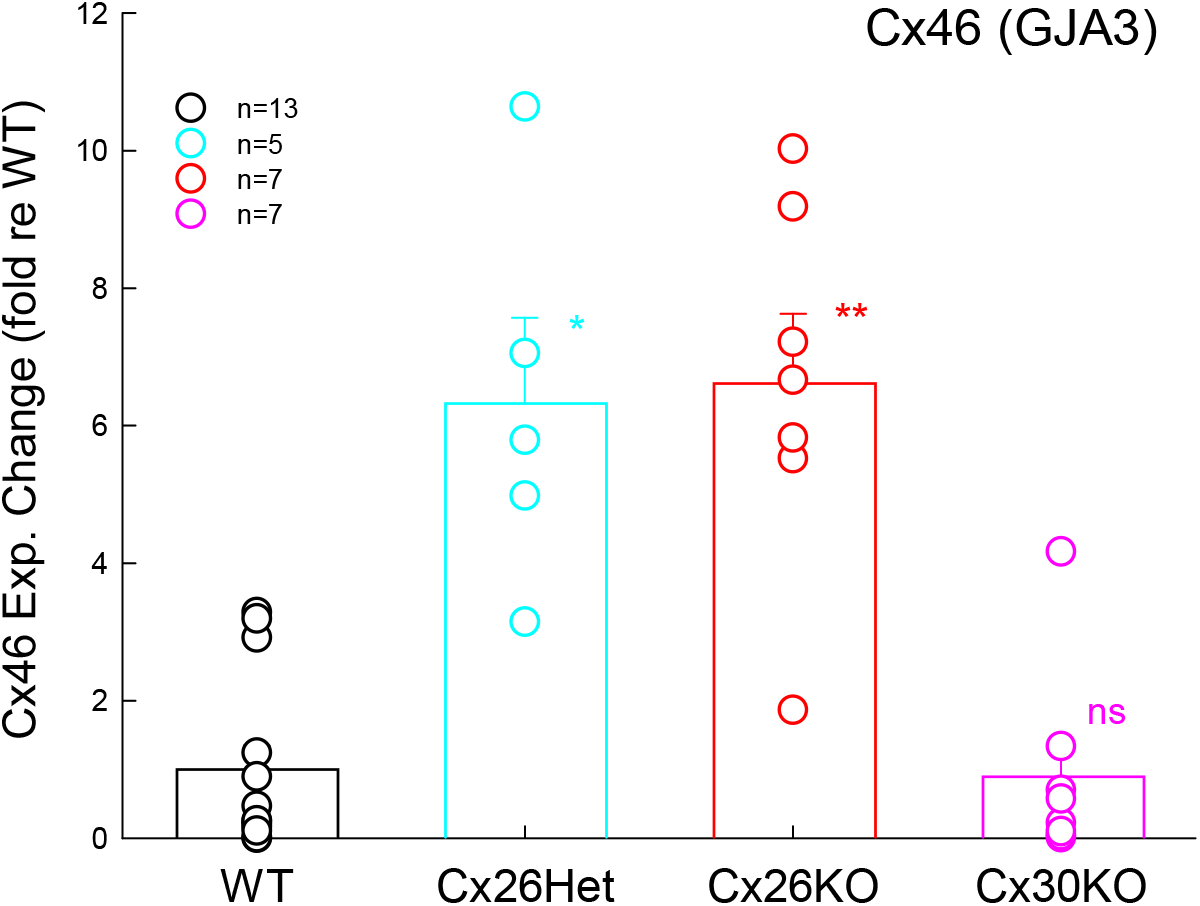
Significant increase of Cx46 expression at the transcriptional level in the cochlea after Cx26 deficiency but not after Cx30 KO measured by real-time quantitative RT-PCR. Significant increase of Cx46 expression at the transcriptional level in the cochlea after Cx26 deficiency but not after Cx30 KO measured by real-time quantitative RT-PCR. Cx46 expression changes were normalized to its expressing level in WT mice. *: P<0.05, **: P<0.01, and n.s.: no significance, one-way ANOVA with a Bonferroni correction.

### No promotion of Cx46 expression in the cochlea after Cx30 deficiency

Cx30 also predominantly co-expresses with Cx26 in the cochlea (14). However, deletion of Cx30 could not prompt Cx46 expression. RNA-Seq shows that Cx46 in the cochlea was not upregulated after Cx30 KO (Fig. 1B). Also, similar to the RNA-Seq results (Fig. 1B), there was no Cx46 upregulation in the Cx30 KO mouse cochlea in quantitative PCR examination (n=7, P=0.86, one-way ANOVA) (Fig. 2).

### Expression and location of Cx46 in the cochlea after Cx26 deficiency

We further used immunofluorescence staining to examine Cx46 expression in the cochlea. Fig. 3 shows that Cx46 labeling in the WT mouse cochlea was undetectable (Fig. 3A, B, and F). However, in Cx26^+/-^ heterozygous deletion mice, Cx46 labeling occurred in the cochlea (Fig. 3C-E). The apparent Cx46 labeling was visible in the organ of Corti and the stria vascularis (SV) and spiral ligament (SPL) in the cochlear lateral wall (Fig. 3C&D). At the cochlear lateral wall (Fig. 3C&D), Cx46 had intensive labeling between SV and SPL, i.e., at the basal cell layer (13). In the cochlear sensory epithelium with whole-mounting preparation (Fig. 3E&F), Cx46 labeling in Cx26^+/-^ mice occurred in the gap junctional network like Cx26 expression (Fig. 3E). Moreover, the new expressed Cx46 integrated into the gap junctional plaque co-expressed with Cx26 (indicated by white arrows in Fig. 3E). In addition, Cx26 labeling (green) in Fig. 3E appears weak in comparison with WT mice (Fig. 3F), consistent with reduction of Cx26 expression in Cx26^+/-^ mice.

**Fig. 3.**
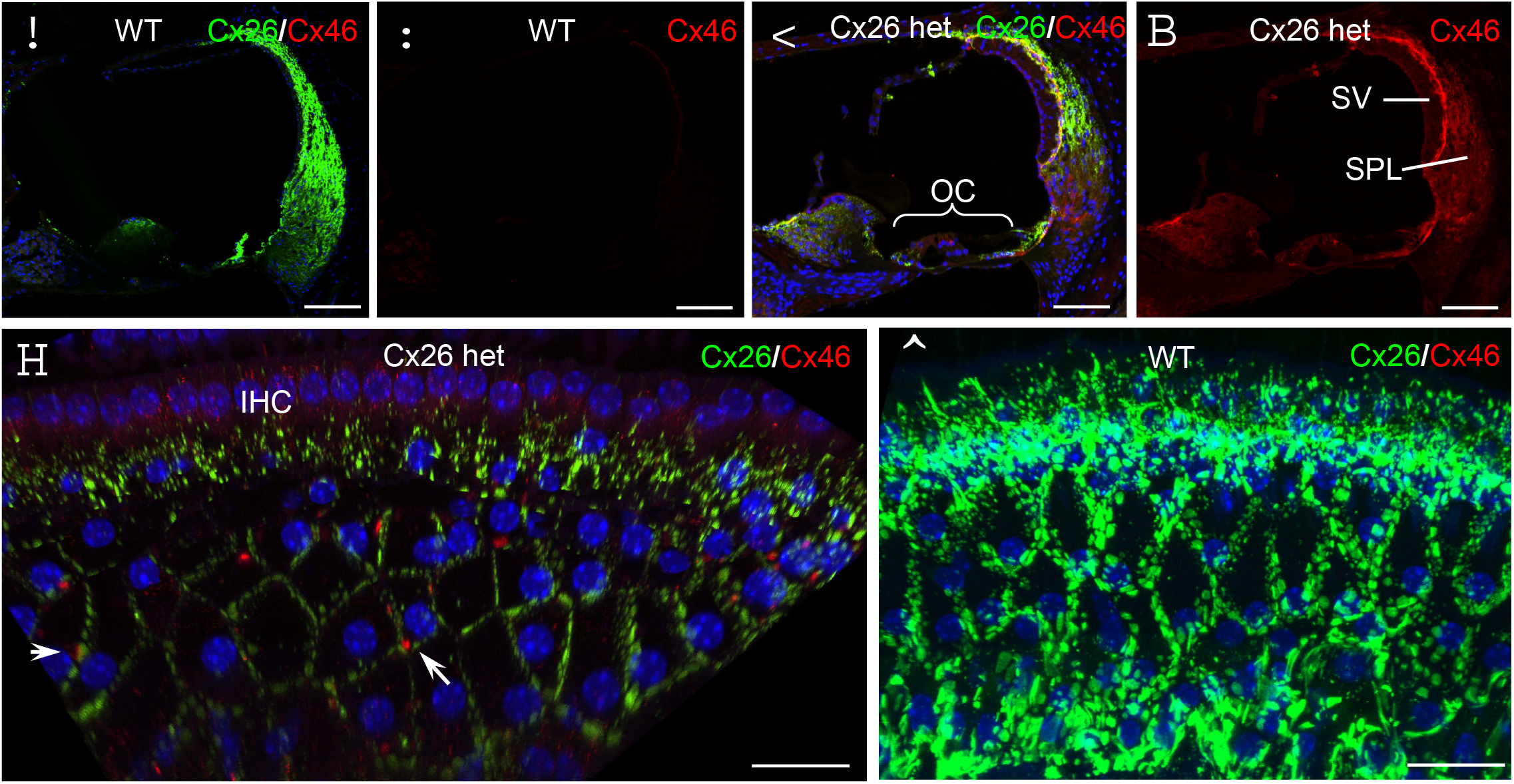
Expression of Cx46 in the cochlea after Cx26 deficiency in immunofluorescent staining. Expression of Cx46 in the cochlea after Cx26 deficiency. **A-B**: Immunofluorescence staining for Cx26 (green) and Cx46 (red) in the cross-section of the WT mouse cochlea. There is no Cx46 labeling (red) is visible in WT mice. **C-D**: Cx46 staining in the Cx26^+/-^ heterozygous deletion mouse cochlea. Labeling for Cx46 (red) is visible at the organ of Corti (OC) and the cochlear lateral wall. SV, stria vascularis; SPL, spiral ligament. **E-F**: Immunofluorescence staining for Cx26 and Cx46 in the cochlea in whole-mounting preparation. White arrows in panel **E** indicate Cx46 labeling occurring at the gap junction network in the cochlear supporting cell area and co-locating with Cx26 at the same gap junctional plaques. However, no Cx46 labeling is detectable in WT mice (panel **F**). Scale bar: 100 µm in **A-D**, 20 µm in **D-F**.

## Discussion

In this study, we found that Cx26 deficiency, but not Cx30 deficiency, promoted Cx46 expression in the cochlea (Figs. 1&2). The newly promoted Cx46 had the same location as Cx26 expression and expressed in the organ of Corti and the cochlear lateral wall (Fig. 3). Moreover, the Cx46 could express in the same gap junctional plaques with Cx26 (Fig. 3E). These data indicate that this promotion of new Cx46 expression is Cx26-specific to compensate Cx26 deficiency.

GJs extensively exist in the inner ear (1) and form two gap junctional networks, i.e., the epithelial cell GJ network in the organ of Corti and the connective tissue GJ network in the cochlear lateral wall (4). However, no GJ and connexin express in the hair cells (14, 19). GJs in the cochlea have many important functions, including participation of cochlear development, maintaining hair cells, participation of regulation of hair cell activity and hearing sensitivity with cochlear efferent system in the organ of Corti, and generation of positive EP in the cochlear lateral wall (4). EP is a driving force and required for generating auditory receptor currents. It has been reported that Cx30 deficiency diminished EP resulting into hearing loss (5). However, in Cx26 KO mice, EP is reduced but not abolished (5). In this study, we found that Cx26 deficiency but not Cx30 deficiency induced promotion of Cx46 expression in the cochlea (Figs. 1-2), especially in the cochlear lateral wall (Fig. 3). This new promoted Cx46 expression may compensate some of Cx26 function letting to part of the EP remaining in Cx26 deficient mice.

Cx46 normally expresses in the eye but not in the ear (15, 16). Different from Cx30, the channel properties of Cx46 are similar to Cx26, having small channel conductance and permeable to both anionic and cationic ions and small molecules (16). This suggests that Cx46 is capable of replacing Cx26 function. Recently, gene therapy has dramatically advanced in treatment of many genetic diseases. Cx26 mutations cause >50% of non-syndromic hearing loss, which is the highest incidence in genetic diseases (1, 3). However, currently, there is no genetic intervention available. Our study found that Cx26 deficiency can promote new Cx46 expressing at the same location of Cx26 in the cochlea after Cx26 deficiency (Figs. 1-3). This study provides a valuable cue to develop a new genetic approach to treat this common genetic disease.

## Acknowledgement

This work was supported by NIH Grant R01 DC019687 to HBZ.

## Author contributions

HBZ designed experiments. TYZ, JC, YK, CL, and HBZ performed experiments and analyzed data. HBZ wrote paper. TYZ, JC, YK, CL, and HBZ reviewed the paper.

## Data and materials availability

All data needed to evaluate the conclusions in the paper are present in the paper.

## Title of new section

Declaration of generative AI and AI-assisted technologies in the writing process.

